# Characterizing Compounds Targeting Colorectal Cancer Derived From Monastrol Using High-Through Screening of an Extensive Combinatorial Library

**DOI:** 10.1101/2025.08.07.667829

**Authors:** Alejandro Rodríguez-Martínez, Lucía Giraldo-Ruiz, Maria C. Ramos, Irene Luque, Diogo Ribeiro, Fátima Postigo-Corrales, Begoña Alburquerque-González, Silvia Montoro-García, Ana Belén Arroyo-Rodríguez, Pablo Conesa-Zamora, Ana María Hurtado, Ginés Luengo-Gil, Horacio Pérez-Sánchez

## Abstract

**Background:** Cancer remains a critical global health concern. Among its various forms, colorectal cancer (CRC) stands out due to its high prevalence and mortality rates, emphasizing the urgent need for novel therapeutic agents to enhance treatment efficacy and prolong patient survival. Monastrol, an antimitotic compound known to bind kinesin Eg5, is employed in some cancer therapies. Recent studies have revealed that monastrol also interacts with fascin, a protein implicated in tumor aggressiveness and metastasis, thereby disrupting microtubule dynamics and actin bundling, ultimately impairing cell migration.

**Methods:** In this work, we developed a workflow to identify fascin-binding compounds based on a monastrol-derived pharmacophore model, integrating *in silico* predictions with *in vitro* validation. We performed ligand-based virtual screening using a pharmacophore model constructed from monastrol, applied to a high-throughput screening (HTS) library of 1.6 million compounds. The top-ranking candidates from the virtual screening were subsequently subjected to physicochemical characterization and cellular assays.

**Results:** Two compounds (designated Z118298144 and Z17544625) were identified that exhibited strong binding to fascin and inhibited actin bundling in physicochemical assays. Furthermore, cellular experiments demonstrated that both compounds reduced proliferation and impaired migration of CRC cells at micromolar concentrations.

**Conclusions:** We established an optimized pipeline combining virtual screening with experimental validation to efficiently identify fascin inhibitors. Using this approach, we discovered two promising compounds with anticancer activity in CRC cell cultures. Moreover, the protocol has been successfully adapted for application to additional cancer-related targets, expanding its potential utility in drug discovery.

**Graphical Abstract:** 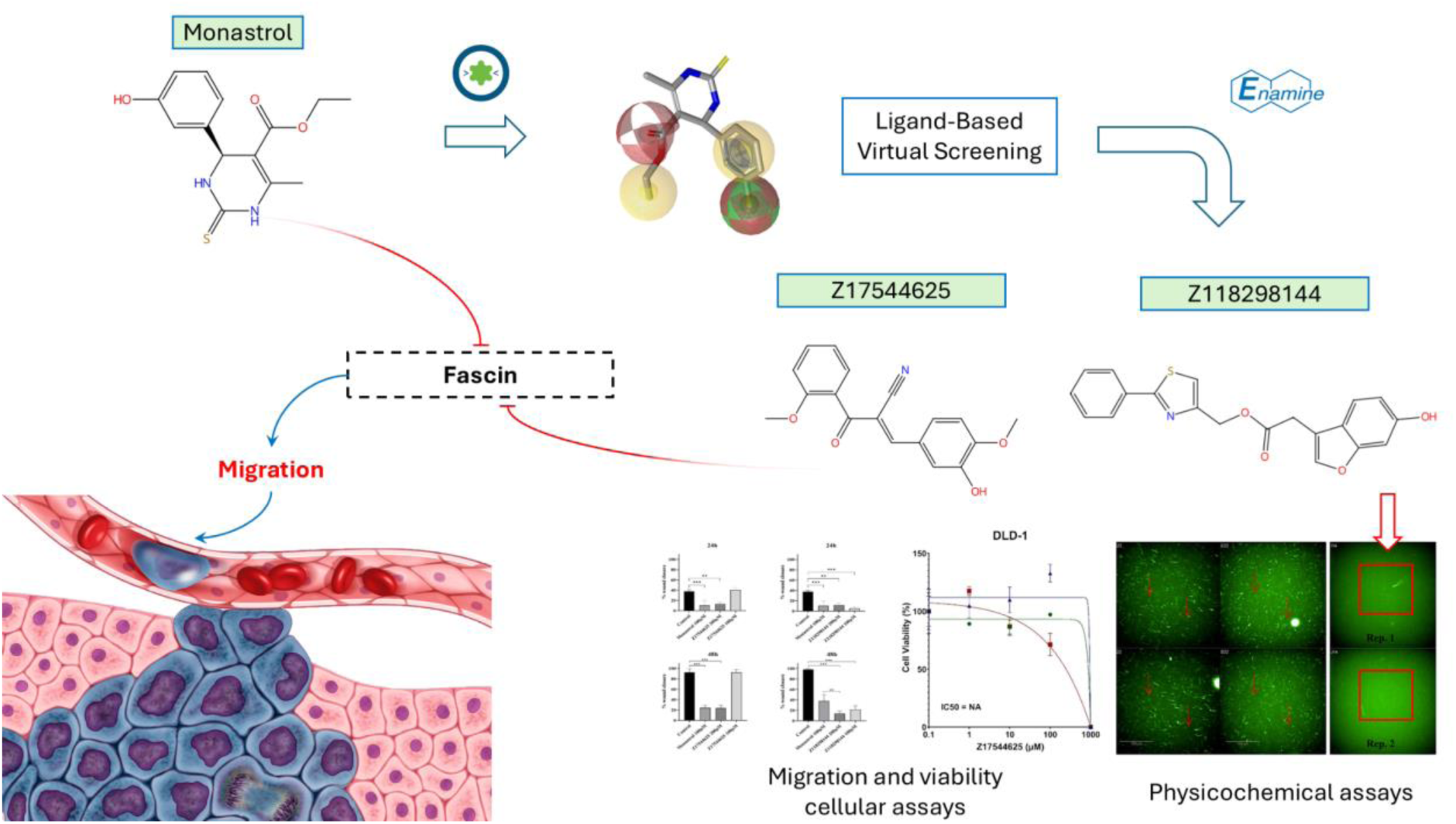

## Introduction

Cancer still poses one of the most significant challenges in healthcare. It is characterized by intricate mechanisms that often circumvent existing treatments, necessitating innovative therapeutic strategies^1^. Over 90% of cancer mortality is due to tumor metastasis, a process underpinned by the capacity of tumoral cells for migration and invasion. This process requires substantial actin-cytoskeleton reorganization to form structures such as filopodia, lamellipodia, and invadopodia, which are crucial for cancer cell mobility^2^.

Eg5 is a kinesin-5-family protein whose high expression levels correlate with poor prognosis in various cancers, including breast cancer, renal cell cancer, and gastric adenocarcinoma^3–5^. Suppression of this protein triggers activation of the spindle assembly checkpoint, resulting in mitotic arrest at the G2/M transition and ultimately inducing apoptotic cell death. Fascin is a key protein in the bundling of actin filaments (F-actin) process and plays a crucial role in migration and invasion processes^6,7^. This protein is significantly upregulated in various human cancers, such as serrated colorectal adenocarcinoma (SAC)^7–9^ and triple-negative breast cancer, thereby increasing the mortality and metastasis risk^10,11^. Monastrol is a small molecule that is recognized as an anticancer agent that arrests mitotic progression by targeting Eg5^12^. Recently, this molecule has also been demonstrated to bind to fascin protein, binding directly and externally to the actin-binding site^13^.

In recent years, *in silico* tools have proven to be efficient in identifying compounds for drug repurposing or even new compounds that act in the desired context. However, in most cases, the search is limited to known libraries of compounds with known drugs or compounds with clinical trials and approval by the FDA. In addition to this type of compound collection, combinatorial chemistry could be useful for investigating new drugs in different contexts. Therefore, we conducted a computational screening of the Enamine HTS chemical library, a collection of 1.368.754 molecules^14^ obtained by combinatorial chemistry and based on the monastrol chemical structure. The compounds with the best results in this screening were tested using *in vitro* physicochemical and cellular assays to assess their effectiveness in halting cancer cell migration and invasion.

## Methods

### Virtual Screening Ligand-Based

#### Generation of HTS Enamine library

The HTS compound library from Enamine, comprising 1,368,049 compounds with diverse chemical characteristics, was utilized for Ligand-Based Virtual Screening calculations. The library was partitioned into 217 files to optimize and parallelize the computational process. Using the LigandScout ‘idbgen’ command, available under its license and executed via the command-line interface, we generated a .ldb format file for each segmented library file.

#### Pharmacophore model generation

In this study, we developed a pharmacophore model for monastrol. The SDF file containing the chemical structure of the inhibitor was sourced from PubChem^15^. This file was uploaded to the LigandScout graphical user interface (GUI)^16^, where the pharmacophore model was generated. Non-essential pharmacophore features that were irrelevant to fascin binding or inhibition were excluded from the model. Additionally, exclusion spheres were added to the filter compounds based on their size and shape. The final refined model was saved in a PMZ format for subsequent screening against the Enamine library prepared in the previous step.

#### Ligand-based Virtual Screening by Ligand Scout

We conducted a comprehensive virtual screening using Metascreener, an in-house software developed by our research group (available at https://github.com/bio-hpc/metascreener). The pharmacophore model was screened against the entire Enamine library, with the maximum number of features omitted (-a, --allow_omit) set to 0, ensuring that all compounds fully matched the pharmacophore features of the G2 model. A post-filtering step was implemented to refine the screening results based on the pharmacophore relative similarity threshold (α = 0.98), as well as economic feasibility and compound availability. As a result, 12 available compounds were selected and ordered from the Enamine shop for subsequent in vitro experiments.

### Physical Chemistry

#### Differential Scanning Fluorimetry

Fascin was assayed at a concentration of 1 μM against 79 compounds in a 384-well plate, with a final assay volume of 10 μL. Using an Echo® 650 Acoustic Liquid Handler (Labcyte, Cat# LP-000061), 400 nL of each compound, prepared from an initial stock of 10 mM in 100% DMSO, was dispensed to achieve a final compound concentration of 400 μM and 4% DMSO. Each assay was performed in quadruplicate to ensure statistically robust measurements.

Two controls comprising 1 μM native fascin were included in the assay: a negative control in 4% DMSO and a positive control for inhibitory activity with 400 μM BDP13176 (MedChemExpress, Cat# HY-119739) in 4% DMSO. Two columns per assay plate were allocated to the controls, with 32 replicates each.

The thermal stability profile of fascin was evaluated using the CFX384 Touch Real-Time PCR Detection System (BioRad, Cat# 1855485) across a temperature range of 25 to 95°C, with a gradient of 1°C/min. The melting temperature (Tm) and Tm shifts induced by compound binding were determined using HTSDSF Explorer software by fitting the fluorescence data to a Boltzmann sigmoidal curve.

The Z’-factor was calculated to measure the robustness and reproducibility of the assay. This Thermofluor assay achieved a Z’-factor greater than 0.75, confirming its high quality.

#### F-actin-Bundling Assay

To visualize F-actin bundling mediated by fascin cross-linking, we implemented an image-based assay following the methodology described by Huang et al. (2018)^17^. F-actin filaments and bundles were labeled using phalloidin conjugated to a fluorescent probe (Alexa Fluor 488-Phalloidin, Thermo Fisher, Cat# A12379), which binds to positive charges of F-actin’s, enabling visualization via fluorescence microscopy. Imaging was performed using the Operetta CLS High Content Analysis System (Revvity, Cat# 8900), and data processing and analysis were conducted using the Harmony™ software (Revvity).

A total of 79 compounds were tested in this assay. Each compound (300 nL), dissolved in 100% DMSO from a 10 mM stock solution, was dispensed using the Echo® 550 Acoustic Liquid Handling system (Beckman Coulter, Cat# 100027). Following this, 15 μL of pure fascin at a concentration of 0.5 μM in buffer (20 mM HEPES, 100 mM NaCl, pH 7.45) was added to a 384-well plate (Corning, Cat# 3677) using an automatic dispenser (Multidrop Combi, Thermo Fisher, Cat# 5840300) and incubated for 30 minutes.

Next, 15 μL of 0.5 μM polymerized actin (prepared in 100 mM KCl, 20 mM Tris-HCl, pH 7.5, 2 mM MgCl₂, 1 mM DTT, and 1 mM ATP; Cytoskeleton Inc., Cat# BK037) was added to achieve a final compound concentration of 100 μM. The plates were then incubated for an additional 30 min. Then, 10 μL Alexa Fluor 488-Phalloidin (diluted 50-fold from 100% methanol stock) was added to stain F-actin, and the samples were incubated in the dark for 1 h. Finally, 25 μL of the resulting solution was transferred to a 384-well poly-D-lysine-coated plate (Corning, Cat# 354663) and incubated for 20 min before imaging.

Each assay plate included 16 wells as negative controls (F-actin without fascin) to confirm that actin bundling was fascin-dependent. Positive controls included the known fascin inhibitor BDP-13176 (Cat# 10009022), which was tested at the same concentration as the compounds. The Z’-Factor, a quality control metric derived from image-defined thickness and texture parameters using the Harmony software, was calculated for each plate. A Z’-Factor greater than 0.5 confirmed the assay’s reliability and robustness.

The results derived from both physical chemistry techniques are provided in the supplementary materials accompanying this publication

### Cell-based assays

#### Cell culture

The human colorectal adenocarcinoma cell lines DLD-1 (CLS Cat# 300220/p23208_DLD-1, RRID: CVCL_0248) and HCT-116 (CLS Cat# 300195/p19841_HCT116.html, RRID: CVCL_0291) were purchased from the American Type Culture Collection (ATCC). The cells were cultured at 37°C in high-glucose Dulbecco’s Modified Eagle’s Medium (DMEM) supplemented with 10% heat-inactivated fetal bovine serum (FBS), 50 U/mL penicillin, and 50 µg/mL streptomycin (Sigma-Aldrich). The cultures were maintained in a 5% CO₂ and 95% humidified atmosphere, and subcultured when the cell confluence reached 90%. Cell line identities were verified using short tandem repeat (STR) profiling.

#### Cell viability assay

Exponentially growing cells were seeded into 96-well plates (Nunc, Roskilde, Denmark) at a density of 1500 cells per well. Each condition was tested in triplicate. After a 24 hours attachment period, serial dilutions of the test compounds (0.1-1000 μM) were added to the wells. Control wells were treated with culture medium containing 0.1% dimethyl sulfoxide (DMSO) (drug carrier). The plates were then incubated for three days at 37°C in a humidified environment with 5% CO₂. Cell viability was determined at the end of the incubation period by using the XTT Cell Viability Assay Kit (Biotium) according to the manufacturer’s protocol. Absorbance was measured at 450 nm with background subtraction at 630 nm using a Spectramax ID3 plate reader (Molecular Devices, VWR).

#### Cell migration assay

The wound-healing assay was conducted using the DLD-1 and HCT-116 cell lines. Cells were seeded at 200,000 cells per well in 35-mm dishes equipped with culture inserts (Ibidi). The assays were performed in triplicate, each including three technical replicates. Four fields per well were imaged to assess migration. After a 24-h incubation period, the inserts were removed to create a defined wound field approximately 500 µm wide. Cell migration was subsequently monitored at 24 and 48 hours in the presence of 10% FBS, following the manufacturer’s protocol. Migration was quantified as the percentage of wound closure using ImageJ,

### Structural Validation

#### Molecular Modeling of fascin structure

Molecular modeling was performed to investigate the interactions between fascin and the predicted compounds. The crystallographic structure of fascin bound to NP-G2-029, a derivative of G2 (PDB code: 6B0T), was used as a starting point.

The structure was preprocessed using the MAESTRO (Schrödinger) software. Chain A of the fascin structure was extracted, and hydrogens and charges were added to complete the structure. The processed structure was then saved in the mol2 format. A parallel protocol was carried out using AutoDockTools software, where similar preprocessing steps were performed, including the addition of AD-specific atom types. In this case, the structure was saved in the pdbqt format.

The predicted ligands were converted into mol2 and pdbqt formats using ChemAxon’s MolConvert tools and AutoDockTools. Hydrogens and their corresponding charges were added to ensure proper molecular representation for the following docking studies.

#### Blind Docking calculations

Blind docking calculations were conducted for two compounds, Z17544625 and Z118298144, which were experimentally shown to bind fascin and reduce cancer cell migration and viability. The crystallographic structure of fascin (PDB code: 6B0T) was used to explore the potential binding sites of these compounds.

Two different docking algorithms, AutoDock Vina and Lead Finder, were employed to perform docking calculations. Both methods were performed to enhance the reliability and coverage of potential binding sites. After completing the docking simulations, a consensus approach was applied, which combined the results from both algorithms to determine the average binding pose for each compound.

#### Molecular Dynamics simulations

Molecular Dynamics (MD) simulations were performed for 100 ns on the fascin structure (PDB code: 6B0T) using the poses with the best docking scores from the Blind Docking (BD) calculations. These poses corresponded to the known binding sites of fascin: actin-binding site 1, actin-binding site 2, and the tubulin-interacting site. The ligand topologies for each pose were generated using ACPYPE^18,19^, and were supported by an automated preprocessing script.

MD simulations were conducted with GROMACS 2022.3^20^ on the Picasso server, utilizing GPU acceleration (NVIDIA A100-SXM4-40GB) and 4 GB of RAM. The following steps were performed:

1. Protein Topology Generation: The gmx pdb2gmx command generated the protein topology, specifying AMBER99SB as the force field.
2. System Preparation: A simulation box was defined, the system was solvated, and ions were added to neutralize the charge.
3. Energy Minimization: A 2000 ps energy minimization stage was performed to remove steric clashes and stabilize the system.
4. Equilibration Stages:

- A single NVT equilibration stage was conducted for 50,000 ps.
- Five consecutive NPT equilibration stages were performed, each lasting 50,000 ps.
5. Production Dynamics: The final MD simulation was run for 100 ns.

The results of the MD simulations were analyzed using ASGARD^21^, an in-house tool developed by our research group. The analysis focused on calculating non-bonded interactions and hydrogen bonds involving the final ligand and evaluating the stability of the protein-ligand complex. Furthermore, the fascin flexibility and dynamic behavior were examined to understand the impact of ligand interactions at the binding site. Input files were prepared accordingly, and the tool generated the necessary plots and raw data for detailed interpretation.

### Drug Feasibility

#### ADME property predictions

Drug-like properties were predicted using the QikProp module of the MAESTRO Schrödinger suite^22^. Calculations were performed for the best-known fascin inhibitors: imipramine, migrastatin, raltegravir, NP-G2-044, G2, BDP-13176, and monastrol, as well as the predicted compounds Z118298144, and Z17544625. The results were saved in a CSV file, and a comparative table was generated to analyze the data.

#### Toxicology profile predictions

Ligand-based screening was conducted using models derived from Z118298144 and Z17544625 against the DrugBank database to identify potential alternative targets of these compounds.

## Results

### Pharmacophore screening

We obtained 1634 hits from the monastrol pharmacophore model. A threshold of Relative-Pharmacophore score=0.98 was chosen to filter most of the compounds obtained by the calculation. Finally, the compounds Z56754204, Z1316642426, Z1177947062, Z57012338, Z930944046, Z44416195, Z17544625, Z788127954, Z1673093246, Z982131356, Z56977400, and Z118298144 were selected for the in vitro assays due to their availability and price for the supplier. Figures 1 and 2 show the 2D and 3D pharmacophore models representation for the two compounds with the best values obtained by physicochemical and cellular experiments (Z17544625 and Z118298144).

**Figure 1.**
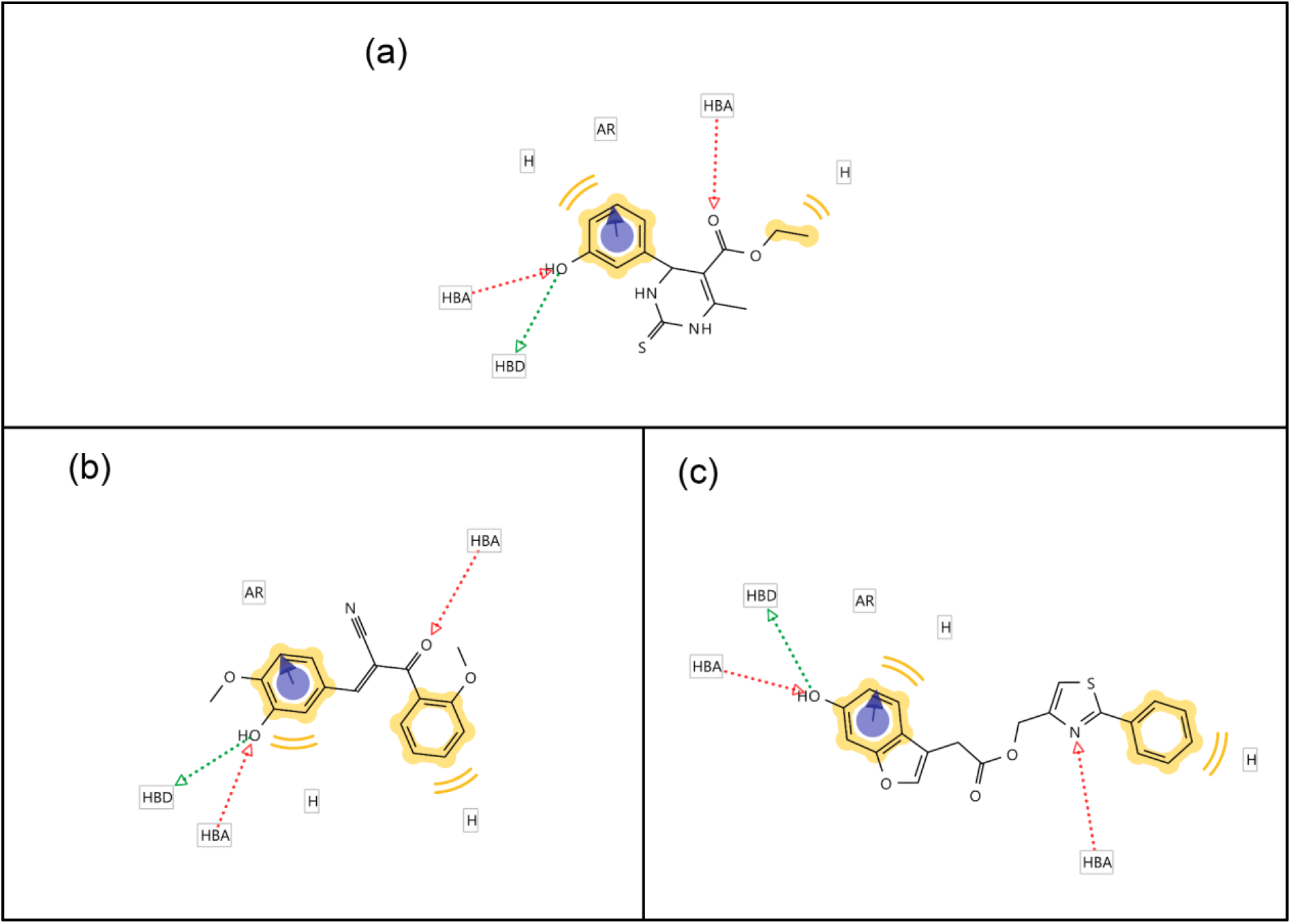
Comparison between the chemical structure and the pharmacophore models of the monastrol (a) and the compounds obtained by Virtual Screening from Enamine library, Z118298144 (b), and Z17544625 (c).

**Figure 2.**
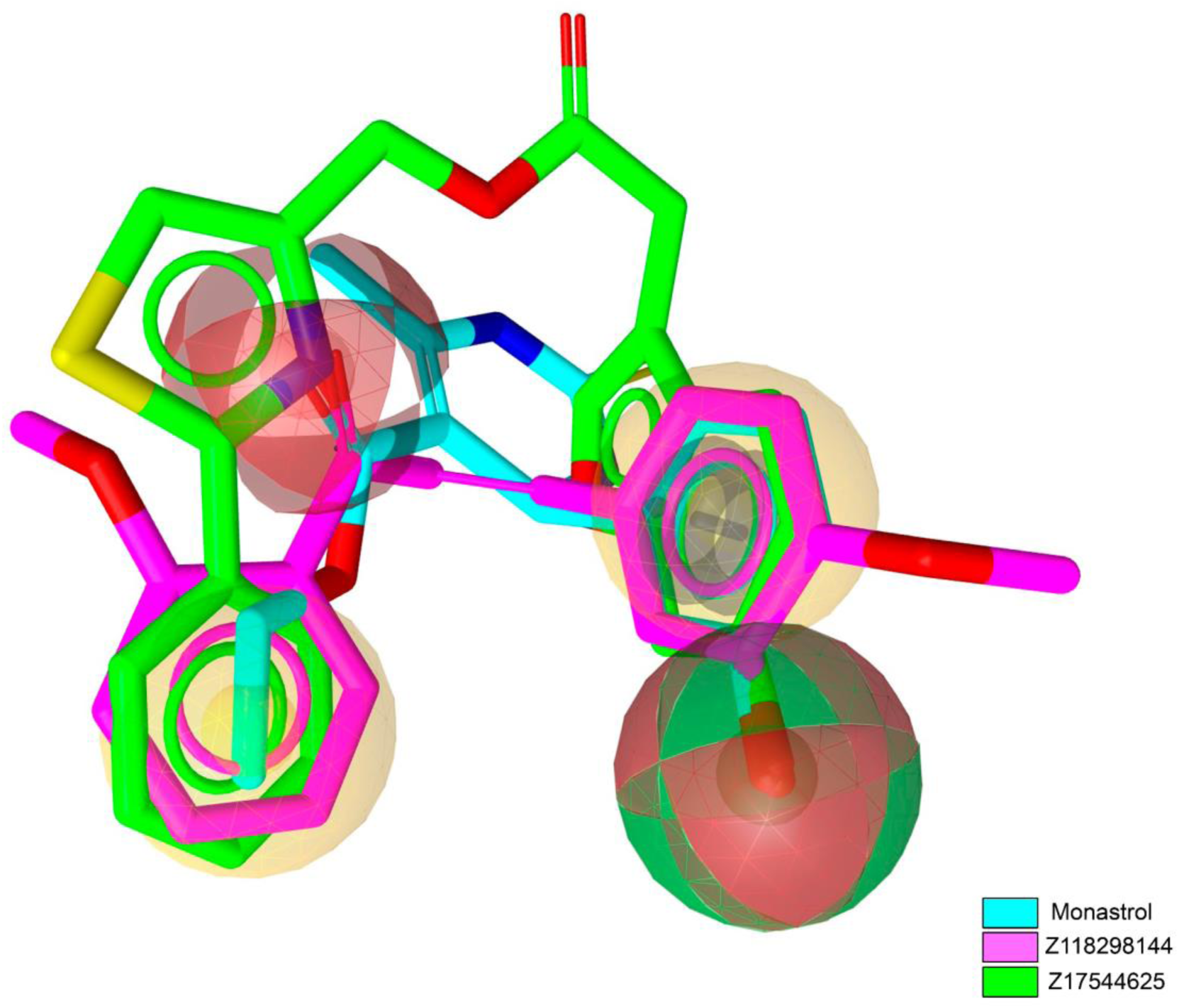
Comparison between the tridimensional chemical structure of monastrol (cyan) and Enamine compounds obtained by Ligand-Based Virtual Screening, Z118298144 (magenta) and Z17544625 (green).

### Viability of cancer cells

Viability assays were performed on DLD-1 and HCT-116 cells to determine the working concentrations of the drugs (Figure 3). These two CRC cell lines exhibit differential endogenous fascin expression levels, with HCT-116 showing higher and DLD-1 lower expression, as previously verified^23,24^

Overall, HCT-116 was more sensitive than DLD-1 in all tested drugs, with Z118298144 exhibiting higher cytotoxicity than monastrol in HCT-116. Furthermore, the obtained IC50 value of monastrol corroborates previously reported findings^13^. Given the differential fascin expression between the cell lines, a comparative evaluation of potential fascin-dependent drug responses was conducted, observing a lower sensitivity in DLD-1 cells that is consistent with their low expression of fascin, as opposed to a superior efficacy in HCT-116.

The approximate working concentrations of monastrol, Z17544625, and Z118298144 were set for subsequent *in vitro* studies at 40 and 100 µM for HCT-116 and DLD-1 cells, respectively. These concentrations were set below the determined IC50 values to minimize cytotoxic effects, enabling reliable assessment of cellular migration in subsequent assays.

**Figure 3.**
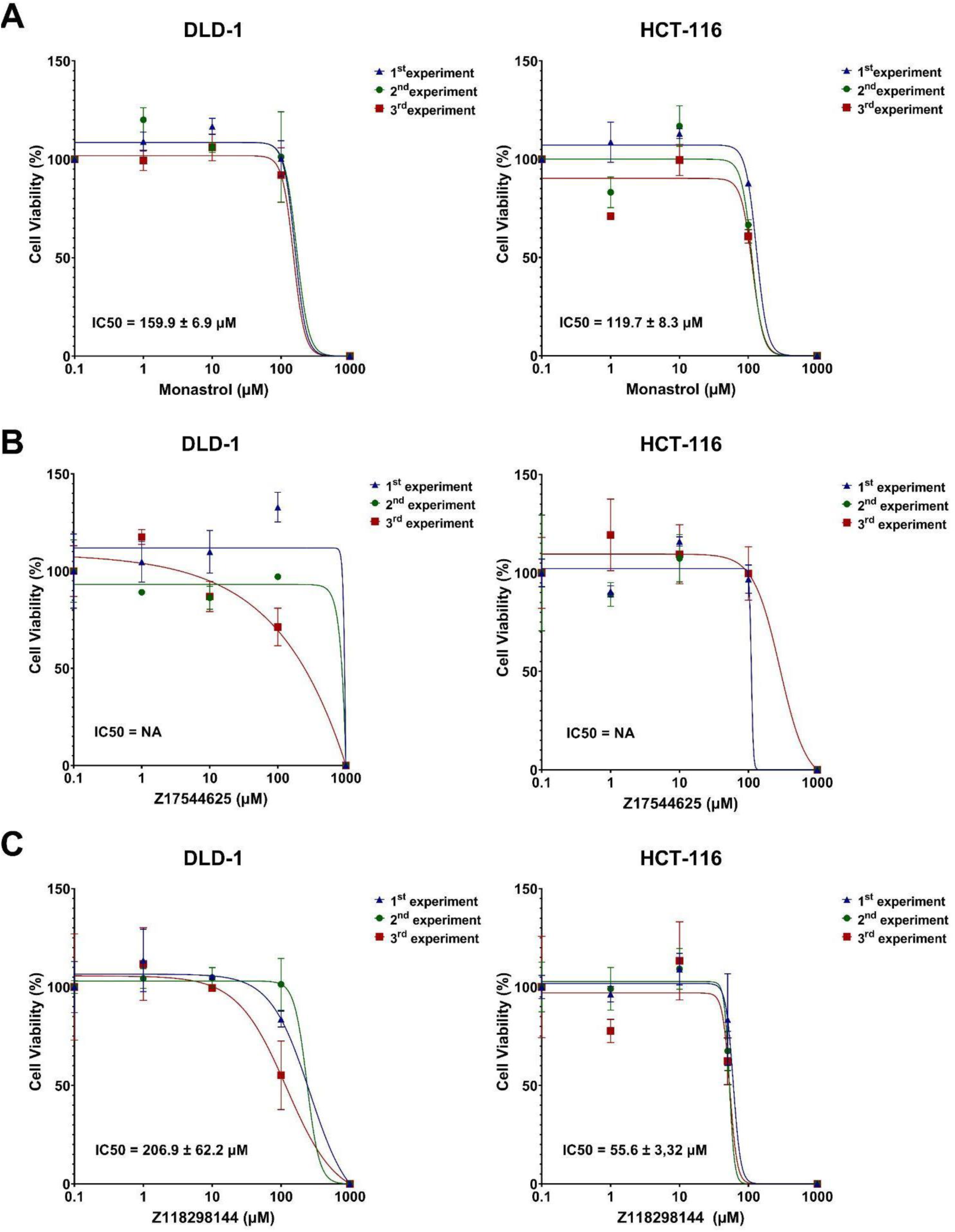
Cell viability assays for DLD-1 and HCT-116 cells were performed in three independent experiments with monastrol (A), Z17544625 (B), and Z118298144 (C). Each line represents the normalized curve fit for an independent experiment. Each point was determined in quintuplicate (mean value + SEM).

### Cancer cells’ migration

To investigate the impact of Z17544625 and Z118298144 on cell migration compared with the monastrol effect, DLD-1 and HCT-116 human CRC cell lines were treated with these compounds at previously established working concentrations, as well as at a higher concentration, to evaluate their effects on cellular migration in a dose-dependent manner. Cell motility was analyzed using a wound-healing assay at 24 and 48 hours. As shown in Figure 4A, all the compounds at 200 µM showed a significant decrease in DLD-1 cell migration, with Z118298144 having the most remarkable effect, even after 48 hours of treatment, and it was significantly more pronounced than monastrol (p < 0.05). Conversely, when the concentration of Z17544625 was reduced to 100 µM, there was no observable impact on cell migration at either of the evaluated timepoints. On HCT-116 cells, the effect of these compounds on migration was only significant for monastrol and Z118298144 (Figure 4 B), with the latter having a similar effect to monastrol at the higher concentration. Notably, Z118298144 exhibited greater cytotoxicity compared to monastrol, which likely contributed to its pronounced inhibitory effect on cell migration at the higher concentration.

**Figure 4.**
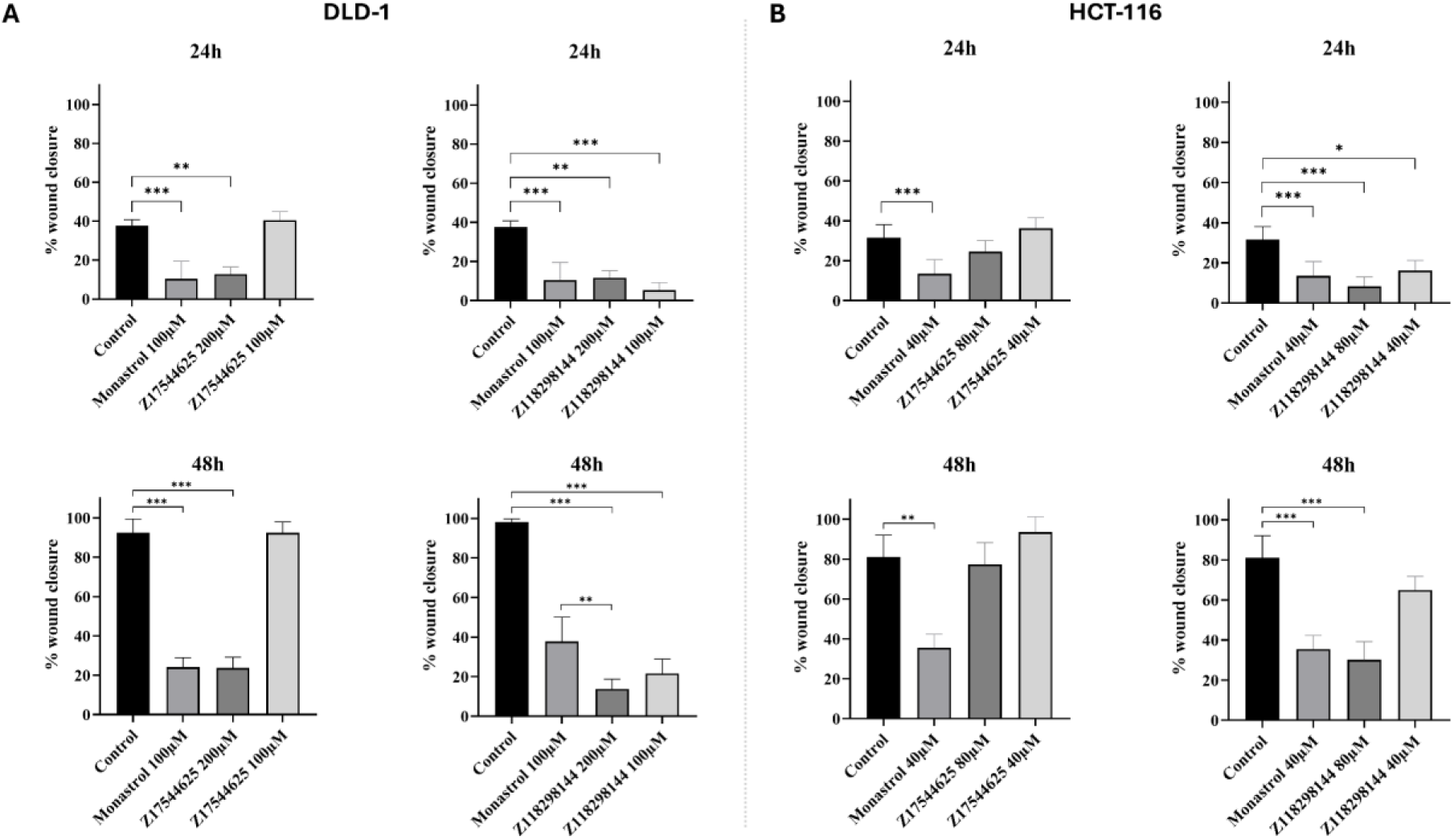
Wound healing assays for DLD-1 (A) and HCT-116 (B) colorectal cancer cells at 24 and 48h using monastrol, Z17544625, and Z118298144.

### Blind Docking detected binding in the Actin-Binding Site 1 and Tubulin Interacting Site for both compounds

Blind Docking calculations using AutoDock Vina and LeadFinder algorithms were performed for the two compounds with the best results (Z118298144 and Z175446245) in the experimental assays against the 6b0t fascin structure to compare with the Blind Docking obtained for monastrol^13^. We found poses in the known binding sites for monastrol: actin-binding site 1, tubulin-interaction site (Figure 3), and actin-binding site 2 (Figure 4).

First, we identified several possible clusters where compounds bind to some residues within the actin-binding site 1 (Table 1)^25^. Z118298144 shows poses interacting with the actin-binding site 1 in clusters 5 and 8 using Lead Finder AutoDock Vina, respectively. Several key residues interact with the ligand (Ser38, Ser39, Lys41, and Lys43). Regarding the Z17544625 compound, we obtained similar results to Z118298144 with cluster 13 for LeadFinder results, and cluster 3 and 10 for AutoDock Vina and interacting with Ala26, Phe29, Phe31, Gly30, Asn34, Ser38, Ser39, and Lys41, known actin site 1 key residues^25^.

**Figure 3.**
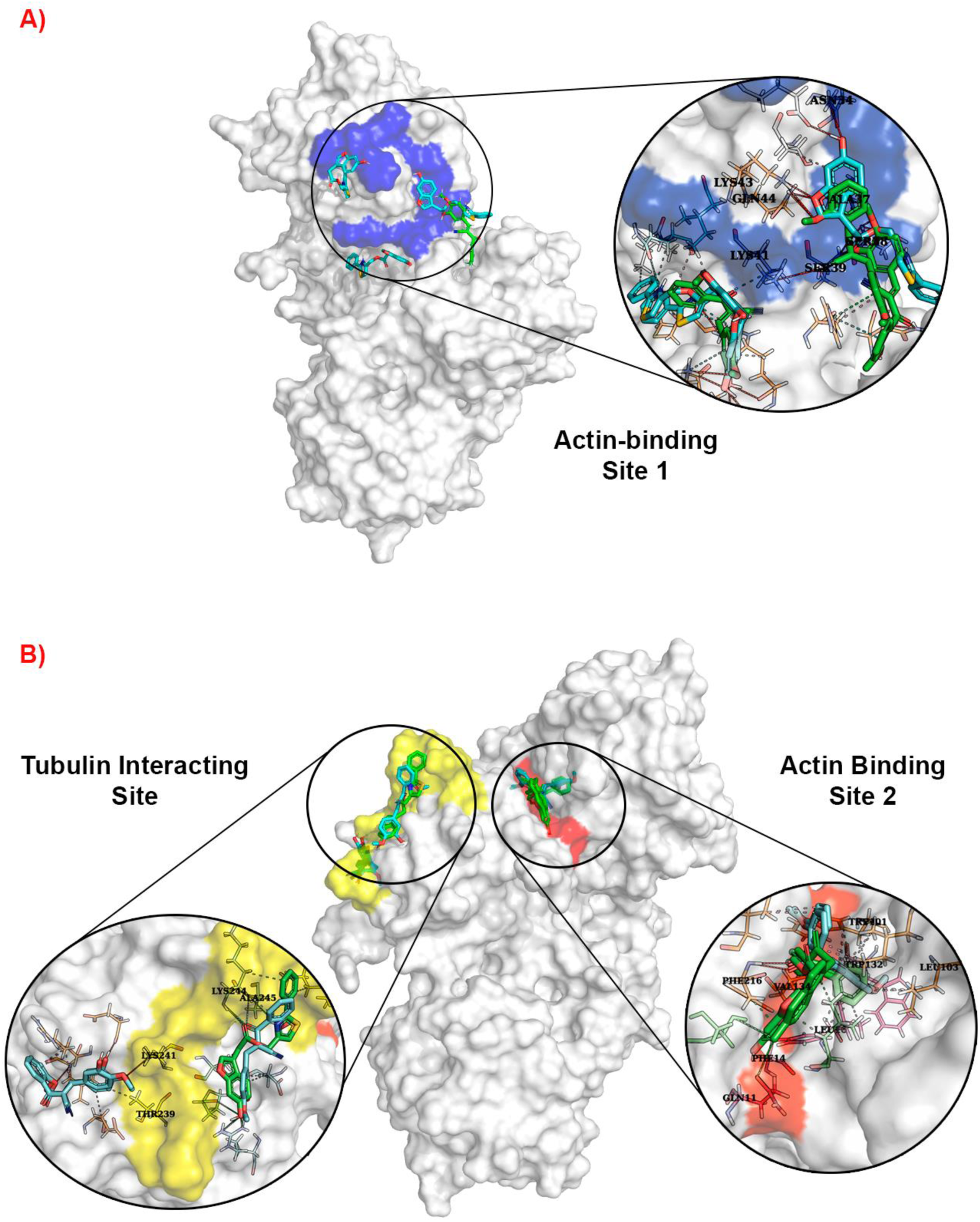
Poses obtained by Blind Docking calculations for Z118298144 and Z17544625 against the fascin protein (6b0t). Green represents the poses obtained for Z118298144 by AutoDock Vina and Lead Finder algorithms. Cian represents the poses obtained for Z17544625. The red and yellow surfaces indicate the actin-binding site 1, the tubulin-interacting site of fascin, respectively. Figure 3.A) shows the poses obtained for the actin-binding site Figure 3. B) shows the poses obtained for the actin-binding site 1 and tubulin-interacting site.

**Table 1.** Interactions detected for compounds Z118298144 and Z17544625 in the fascin Actin-Binding Site 1 by AutoDock Vina and Lead Finder Docking calculations.

|  | Compound | Z118298144 | Z17544625 |
| --- | --- | --- | --- |
| LF | Score (kcal/mol) | **-6.64** | **-5.94** |
|  | Hydrophobic Interactions (Å) | Asn407 (3.97) | **Ala26 (3.53), Phe29 (3.92), Phe29(3.30), Phe31 (3.30),**<br>Leu66 (3.54, 4.00), Arg68 (3.69) |
|  | Hydrogen Bonds (Å) | **Ser38 (2.89), Ser39 (2.83, 3.02),** Asn407 (2.81) | **Gly30 (2.84)** |
|  | Salt Bridges (Å) | - | - |
| | π-Cation Interactions (Å) | His392 (4.09) | - |
| AD | Score (kcal/mol) | **-5.8** | **7.49, 6.42\*** |
|  | Hydrophobic Interactions (Å) | His392 (3.82), Asn407 (3.83) | Ile12 (3.72), Gln13 (3.71), **Lys41 (3.95)**, Ile45 (3.70), Met138 (3.49)<br>Thr25 (3.97), **Ala37 (3.60)** |
|  | Hydrogen Bonds (Å) | **Ser38 (2.84), Ser39 (3.93, 2.85), Lys41 (3.18), Lys43 (3.84),** Gln44 (3.93), Asn407 (3.38) | Gln13 (2.80)<br>Glu27 (2.84), **Asn34 (3.36), Ser38 (3.04, 3.28, 3.28), Ser39 (3.07), Lys43 (2.80),** Gln44 (3.20) |
|  | Salt Bridges (Å) | - | Arg389 (4.22)<br>- |
| | π-Cation Interactions (Å) | His392 (3.94) | - |

\*Two possible poses for that site were found.

The actin-binding site 2 showed the best results, located in the first pose in both compounds and by both algorithms. The docking scores for Z118298144 are -8.14 and -7.4 kcal/mol for LeadFinder and AutoDock Vina, respectively. Regarding the Z175446245 compound, the docking scores were -9.72 kcal/mol and -9.16 kcal/mol calculated by LeadFinder and AutoDock Vina, respectively. Table 2 shows the interactions detected by both methods, and bold residues correspond to the residues belonging to each site^17,26^. Thus, in both cases, several residues participating in each of these domains interacted with the molecules identified by the screening.

**Table 2.** Interactions detected for the compounds Z118298144 and Z17544625 in the fascin Actin-Binding Site 2 by AutoDock Vina and Lead Finder Docking calculations.

|  |  |  |  |
| --- | --- | --- | --- |
|  | Compound | Z118298144 | Z17544625 |
| LF | Score (kcal/mol) | -8.14 | -9.72 |
|  | Hydrophobic Interactions (Å) | Ile93 (3.41, 3.92), Trp101 (3.15), Leu103 (3.48), Val134 (3.22), Glu215 (3.97) | Phe14 (3.59, 3.64), Leu16 (3.69), Leu48 (3.20,3.03), Ile93 (3.59), Trp101 (3.63, 3.84), Leu103 (3.84), Phe216 (3.18) |
|  | Hydrogen Bonds (Å) | Trp101 (2.84), Trp132 (3.97), Phe216 (3.11) | Gln11 (3.09), Gln12 (2.74), Phe216 (4.06), Ser218 (2.69), Gly219 (3.84) |
|  | Salt Bridges (Å) | - | Arg217 (4.51) |
| AD | Score (kcal/mol) | -7.4 | -9.16 |
|  | Hydrophobic Interactions (Å) | Ile93 (3.46,3.34), Trp101 (3.34), Leu103 (3.30), Val134 (3.45), Glu215 (3.77) | Phe14 (3.50), Leu16 (3.92), Leu48 (3.91,3.73), Val60 (3.84), Ile93 (3.81,3.71), Trp101 (3.73, 3.72), Leu103 (3.73) |
|  | Hydrogen Bonds (Å) | Trp101(3.05), Phe216 (3.13) | Gln11(2.89), Ser218 (3.04) |
|  | Salt Bridges (Å) | - | Arg217 (4.10) |

Regarding the tubulin-interacting site (Table 3), for compound Z118298144, both methods found fewer possible poses in both docking softwares. Otherwise, the LeadFinder approximation did not find any cluster with interactions in some of the residues corresponding to the tubulin-interacting site of Z17544625. AutoDock Vina obtained a pose close to the tubulin-interacting site in cluster 13 with -6.17 kcal/mol of docking score. Regarding the key residues, some residues indicated as belonging to the tubulin-interacting site in the literature^26^ were found by both techniques (Thr239, Lys241, Lys244, Ala245, and Lys247).

**Table 3.**
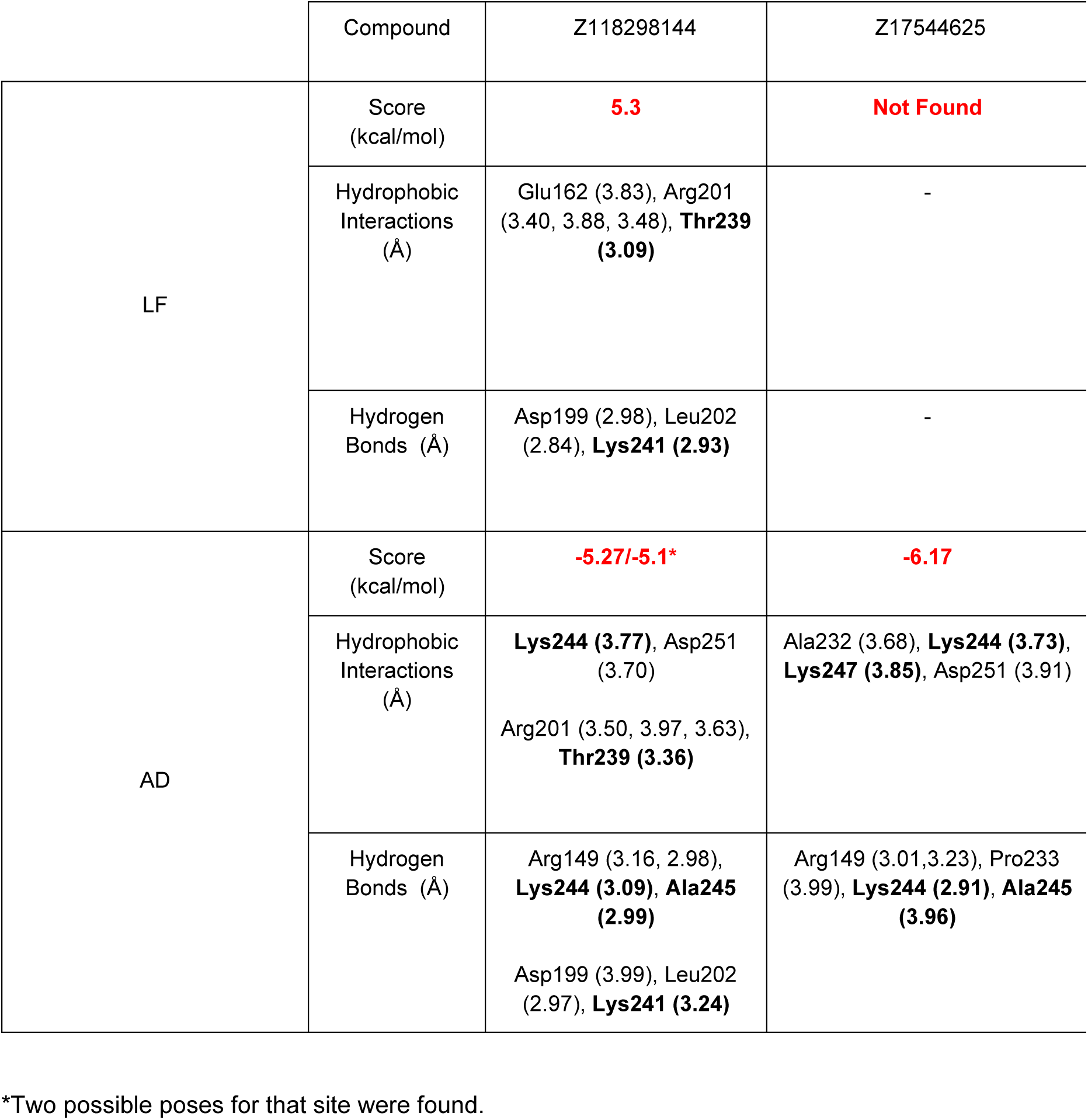
Interactions detected for the compounds Z118298144 and Z17544625 in the fascin Tubulin-Interacting Site by AutoDock Vina and Lead finder Docking calculations.

|  | Compound | Z118298144 | Z17544625 |
| --- | --- | --- | --- |
| LF | Score (kcal/mol) | **5.3** | **Not Found** |
|  | Hydrophobic Interactions (Å) | Glu162 (3.83), Arg201 (3.40, 3.88, 3.48), **Thr239 (3.09)** | - |
|  | Hydrogen Bonds (Å) | Asp199 (2.98), Leu202 (2.84), **Lys241 (2.93)** | - |
| AD | Score (kcal/mol) | **-5.27/-5.1\*** | **-6.17** |
|  | Hydrophobic Interactions (Å) | **Lys244 (3.77)**, Asp251 (3.70)<br>Arg201 (3.50, 3.97, 3.63), **Thr239 (3.36)** | Ala232 (3.68), **Lys244 (3.73)**, **Lys247 (3.85)**, Asp251 (3.91) |
|  | Hydrogen Bonds (Å) | Arg149 (3.16, 2.98), **Lys244 (3.09)**, **Ala245 (2.99)**<br>Asp199 (3.99), Leu202 (2.97), **Lys241 (3.24)** | Arg149 (3.01,3.23), Pro233 (3.99), **Lys244 (2.91)**, **Ala245 (3.96)** |

\*Two possible poses for that site were found.

### Molecular Dynamics simulations verify the binding in our site

We performed three different simulations for each compound, targeting each one to the known monastrol-binding sites (actin-binding sites 1 and 2, and tubulin-interaction site). We carried out MM-PBSA and RMSD analyses to prove the stability of the ligand in the fascin complexes. The poses obtained in the tubulin-interacting site and actin-binding site 1 did not remain in the corresponding binding sites. In these cases, the complexes with fascin showed poor stability, and the compounds left the complex from the first 10-15 ns simulation. In contrast, compounds Z118298144 and Z17544625 were stable in complex with fascin binding to the actin-binding site 2. MD simulation results demonstrated stable binding at the actin-binding site 2 for both compounds, Z118298144 and Z17544625.

According to the data obtained by MD, both compounds had a higher affinity for the actin-binding site 2. The RMSD (Root Mean Square Deviation) of the ligand for both MD (Molecular Dynamics) simulations showed apparent stabilization, remaining in the range of 0.1 to 0.2 nm. Regarding the protein, fascin is known for its high flexibility, displaying fluctuations, especially in complex with Z17544625. However, these fluctuations do not exceed 5 nm.

**Figure X.**
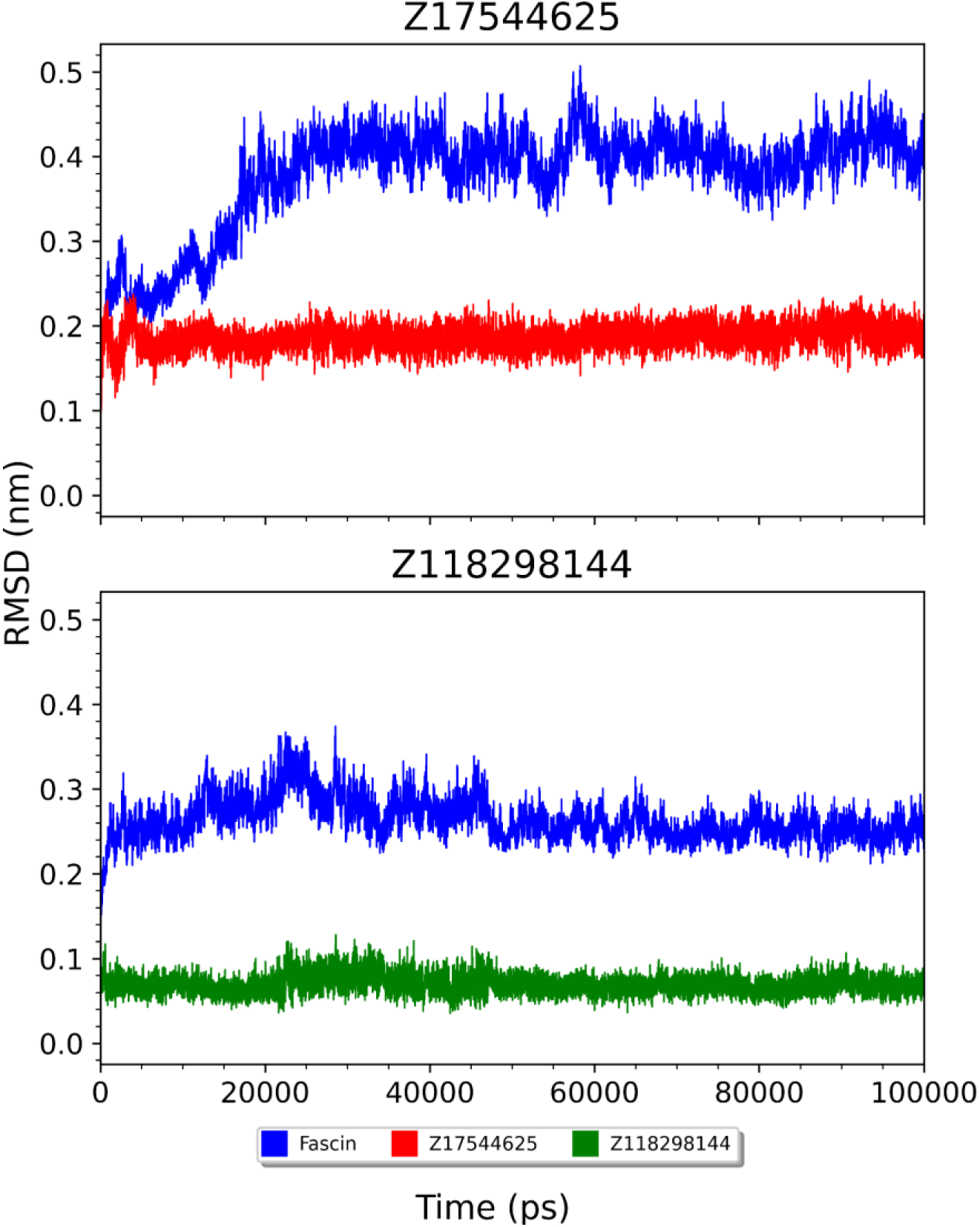
RMSD of fascin and the Z17544 compound for each corresponding MD simulation. The blue color indicates the protein in both cases, the red color indicates the Z17544625 compound in the MD simulation for the complex in the actin-binding site 2, and the green color indicates the Z118298144 compounds in the same case for its simulation.

Finally, MMPBSA demonstrated the stability of this complex binding. As shown in Figure Y, the binding affinities of both complexes are stable between -300 and -350 kJ/mol in the Z17544625 compound and between -200 and -250 kJ/mol for the Z118298144 compound. In addition, there were no strong fluctuations in the energy, which could mean a loss of binding and the formation of the complex. The difference in mean energy during the simulation between both compounds correlated with the data obtained by IC50 assays for *in vitro* assays, showing Z17544625 as a more effective compound.

**Figure Y.**
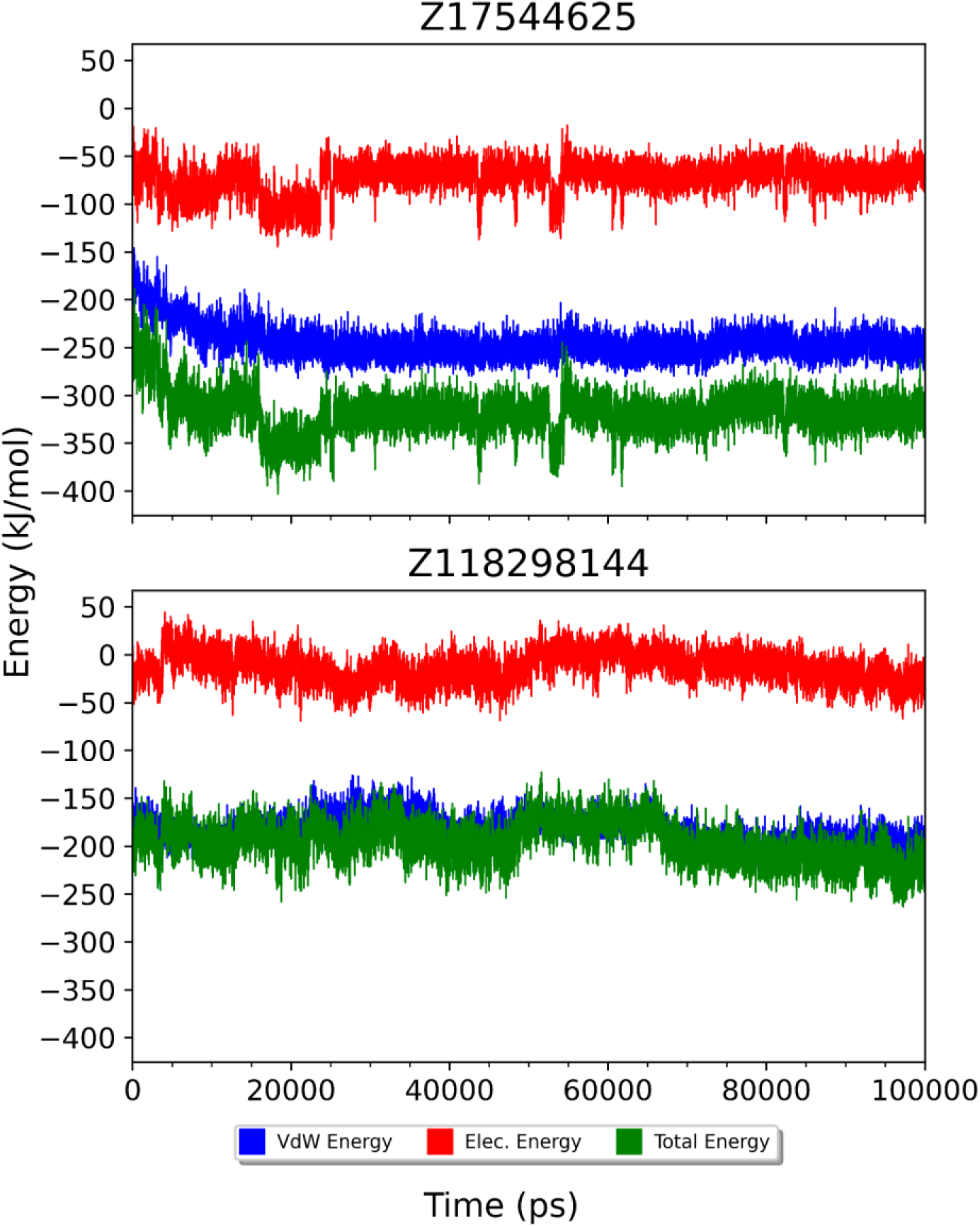
MMPBSA analysis showed the binding energies of the compounds Z17544625 and Z11729144 in the actin-binding site 2 for fascin during the simulation time. The plots indicate the van der Waals (blue), electrostatic (red), and total binding (green) energies .

### Comparing ADMET properties against monastrol

The compounds obtained by the screening had 0 stars in the values of the descriptors determined by QuikProp, similar to monastrol. This rating indicates that all the ADMET-predicted properties of the compounds fall within the optimal range for 95% of the known drugs. Most of the predicted properties displayed similar values for the three compounds analyzed. Table 4 displays the values of the most notable predicted properties. All the values were within the acceptable range for a drug. Additionally, there were no significant differences between the values obtained for the compounds and those obtained for monastrol. These results suggest that the new compounds possess favorable druggability profiles, with ADMET properties closely resembling those of the reference compound, monastrol.

**Table 4.** ADMET properties calculated for monastrol, Z118298144, and Z17544625 by QikProp.

| Molecule | MW<sup>a</sup> | HBD<sup>b</sup> | HBA<sup>c</sup> | Po/w<sup>d</sup> | S<sup>e</sup> | HERG<sup>f</sup> | BB<sup>g</sup> | Khsa<sup>h</sup> | HOA<sup>i</sup> | R5<sup>j</sup> | R3<sup>k</sup> | Jm<sup>l</sup> |
| --- | --- | --- | --- | --- | --- | --- | --- | --- | --- | --- | --- | --- |
| Monastrol | 292.35 | 1.00 | 3.25 | 3.25 | -4.65 | -4.43 | -0.51 | 0.35 | 100 | 0 | 0 | 0.01 |
| Z118298144 | 309.32 | 1.00 | 5.75 | 2.37 | -4.34 | -5.35 | -1.32 | -0.14 | 87.2 | 0 | 0 | 0.04 |
| Z17544625 | 365.40 | 1.00 | 4.75 | 3.51 | -3.48 | -4.24 | -0.34 | 0.26 | 100 | 0 | 0 | 2.6 |

**a:** Molecular Weight. Values within the range: 130.0 - 725.0, **b:** Hydrogen Bond Donor. Values within the range: 0.0 - 6.0, **c:** Hydrogen Bond Acceptor. Values within the range: 2.0 - 20.0, **d:** QP log P for octanol/water. Values within the range: -2.0 - 6.5, **e:** QP log S for aqueous solubility. Values within the range: -6.5 - 0.5, **f:** log IC50 HERG K<sup>+</sup> Channel Blockage: Values within the range: > -5, **g:** QP log BB for brain/blood. Values within the range: -3.0 - 1.2, **h:** QP log K hsa Serum, Protein Binding. Values within the range: -1.5 - 1.5, **i:** Human Oral Absorption in the gastrointestinal tract (%). Values within the range: > 25%, **j:** Lipinski Rule of 5 Violations. Values within the range: < 4, **k:** Jorgensen Rule of 3 Violations. Values within the range: < 3, **l:** Maximum transdermal transport rate (µg/cm<sup>2</sup>/h). Values within the range: 1 - 10.

### Drug-Target Similarity Prediction

Finally, we performed ligand-based calculations for Z118298144 and Z17544625 against the DrugBank compound library to complement ADMET analysis. Thus, some possible alternative drug targets were searched for similarities to other drugs. First, we studied the Z118298144 compound. Here, we observed some interesting targets that obtained high relative pharmacophore scores. First, any possible target that could cause compliance due to the administration of the compounds was identified. The possible targets of interest are Glucose Transporter 1 (GLUT1) and epidermal growth factor receptor (EGFR), two targets studied to block cell proliferation and migration in the context of cancer^27,28^. These targets were identified because of the high similarity of known inhibitors with Z118298144, WZB-117, and Tyrphostin B48, which interact with GLUT1 and EGFR, respectively.

**Table 5.**
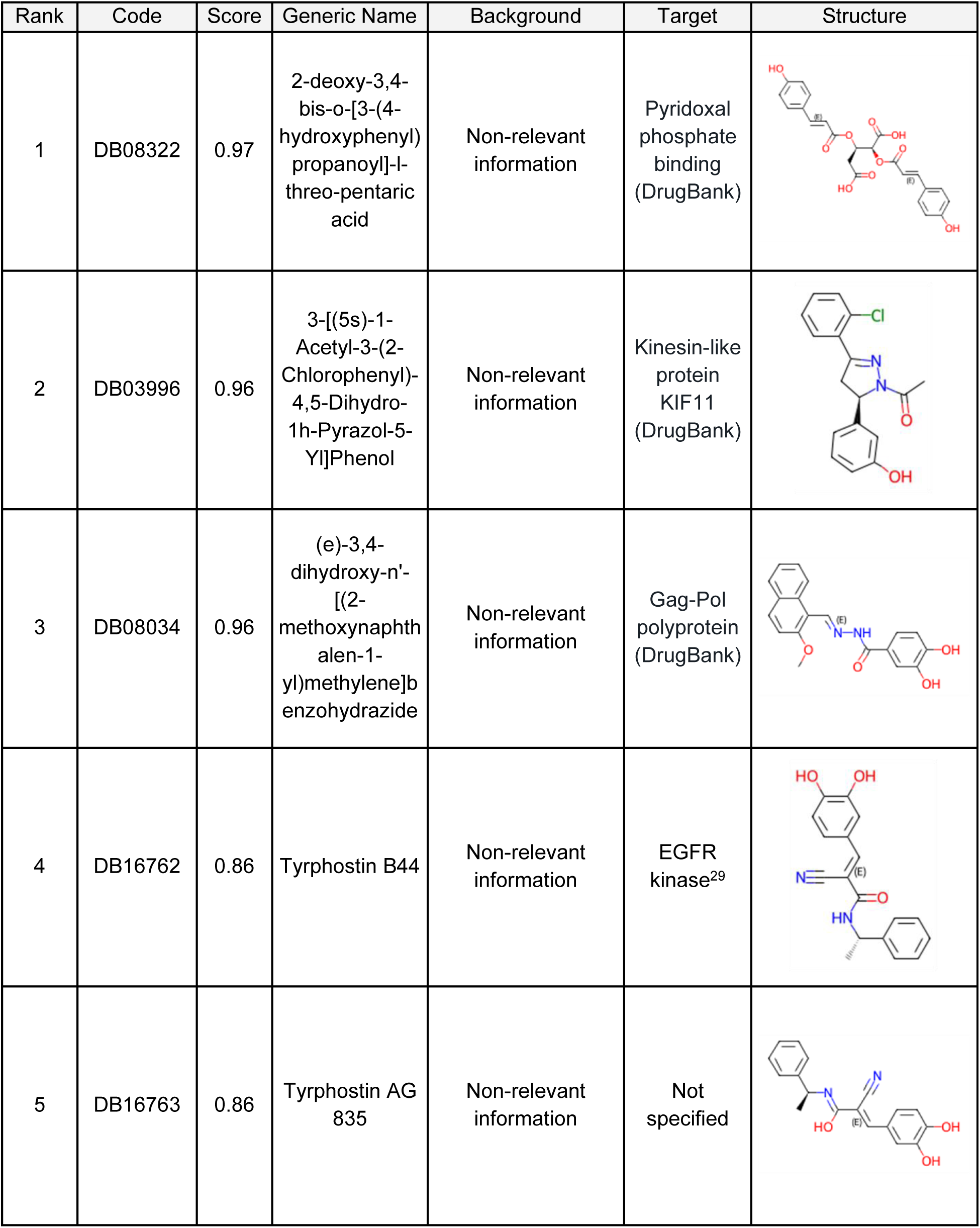
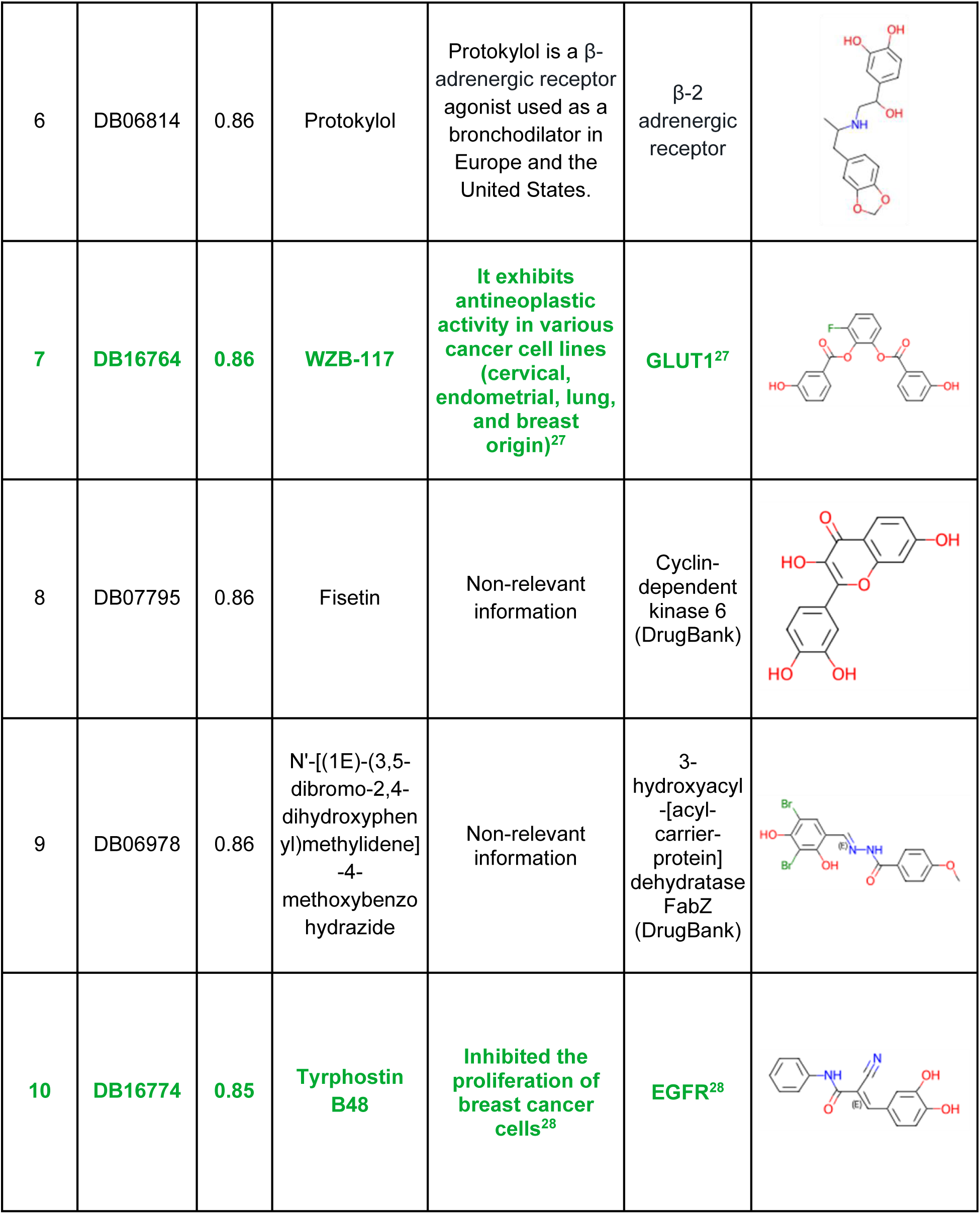
Drug-Target Prediction by Ligand-Based LigandScout of Z118298144.

The study of possible drug targets for the Z17544625 compound also revealed interesting information. As with the Z118298144 compound, any target that could cause adverse effects was found. Tryphostin AG 555 was identified as the second-best hit by pharmacophore similarity for the entire DrugBank library. This compound is related to some cancers, interacting with known cancer-regulating targets, such as tyrosine kinase and EGFR^30^. The other compound was found to be A-674563. This molecule inhibits the CDK2 enzyme, blocking the cell growth in some types of lung cancer^31^.

**Table 6.**
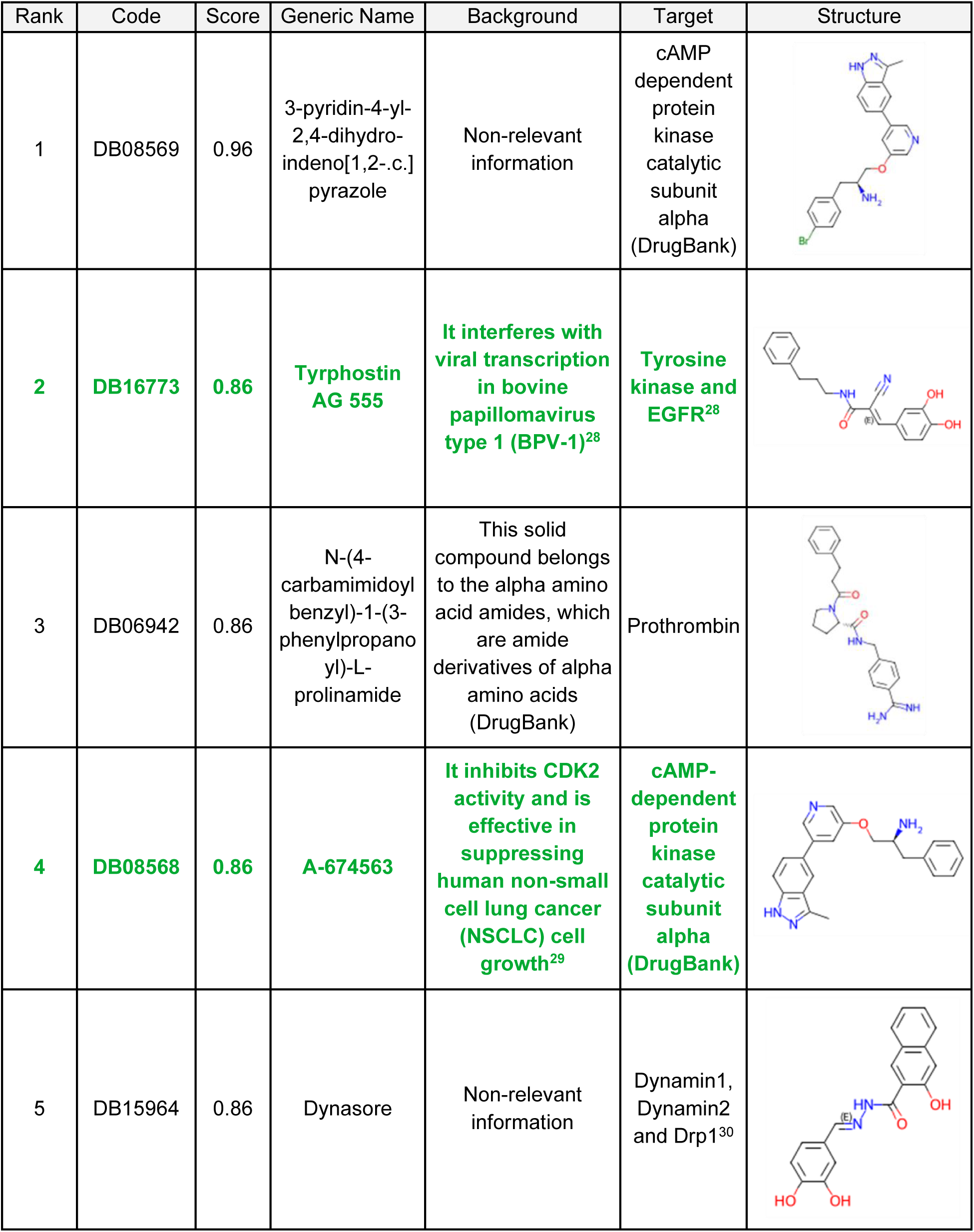
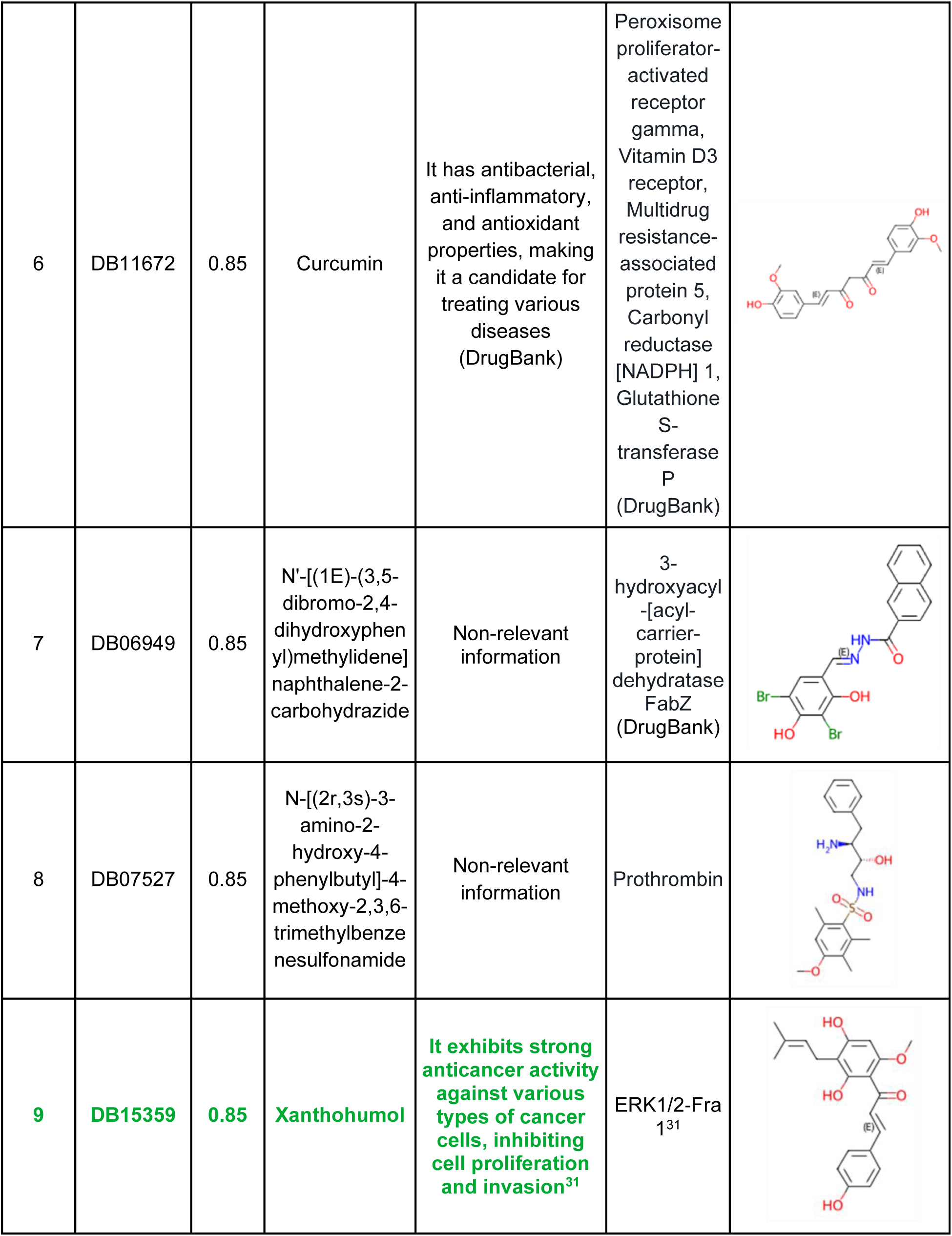
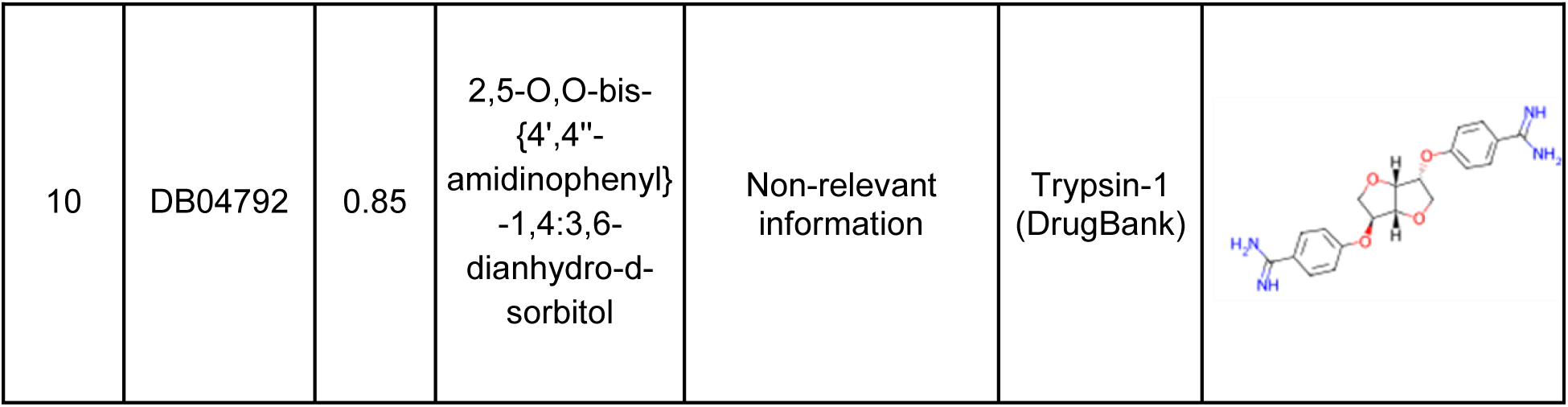
Drug-Target Prediction by Ligand-Based LigandScout of Z17544625.

## Discussion

Monastrol is a small-molecule inhibitor of the mitotic kinesin Eg5, which is essential for maintaining spindle bipolarity during cell division^12^. Recent studies have demonstrated that this molecule also effectively inhibits fascin and blocks migration and metastasis processes^13^. Fascin plays a causative role in tumor migration and invasion and is associated with tumor subtypes with adverse prognosis. This protein is considered a promising therapeutic target and clinical trials with fascin inhibitors are ongoing^34,35^.

Progress in drug discovery in the fascin context is closely linked to screening FDA-approved compound libraries in search of drug repurposing opportunities. This approach identifies existing drugs that may present fewer challenges when advancing clinical trials^36^. However, exploration of increasingly extensive chemical spaces is an underexploited avenue for Drug Discovery.

In this study, we performed virtual screening calculations against a library of over one million compounds, such as Enamine, to identify a compound with activity similar to monastrol binding, such as the fascin protein, and inhibiting tumor migration. The workflow to obtain these possible hits resulted in two compounds with similar activities to monastrol: Z118298144 and Z17544625. *In vitro* assays demonstrated that these compounds also prevented the actin-bundling mediated by fascin and colorectal tumor cell viability and migration, especially Z118298144, whose effect was even more pronounced than that of monastrol.

The *in silico* structure-based analysis showed the stability for both compounds in at least one possible binding site. From 6b0t Blind Docking, we observed that both compounds bind and remain in the known fascin-binding sites for actin and tubulin, which are related to actin-bundling and posterior cancer cell migration. The posterior 100 ns MD simulation for each best pose indicated a stable and strong binding in the actin-binding site 2 for the two compounds.

In addition, the compounds obtained share many drug-like properties, converting these molecules into possible alternative molecules to monastrol and obtaining an excellent value for future rounds of screening to optimize the process and obtain better hits.

In this way, two potential compounds were identified using a screening workflow, including ligand-based virtual screening from the monastrol model, in vitro physicochemical and cellular assays, and Structure-Based computational techniques: Blind Docking and Molecular Dynamics. The ADMET properties and similarity with other drug targets were described to check the drug capability of these compounds, demonstrating a positive evaluation of this methodology, which could help subsequent screening cycles to obtain more effective compounds.

We also introduced a new workflow designed to enhance prediction accuracy of virtual screening for fascin-binding compounds. This workflow integrates several experimental validation methods, including physicochemical analyses and cell culture-based assays. Our results demonstrated that this tool is easy to implement and efficient. The computational pipeline begins with a single input file representing the molecular structure of a known fascin inhibitor monastrol and proceeds through a series of automated steps that enable the identification of top-scoring hits from an HTS chemical library. Notably, two of these hits exhibited enhanced inhibition of actin bundling and improved antiproliferative and antimigratory activities compared with monastrol.

This strategy may serve as a valuable platform for the discovery of novel compounds with higher activity than that of the reference molecule. Through iterative application, this workflow could facilitate the identification of candidates with micromolar and even nanomolar IC50 values and significant potential for cancer treatment. We intend to apply this methodology to other established anticancer targets, thus contributing to the ongoing evolution of drug discovery practices.

Despite its promise, the current workflow has limitations that should be addressed in future studies. For example, it relies solely on pharmacophore similarity for ligand-based hit identification. Incorporating additional ligand-based approaches, such as fingerprint similarity (DRAGON, PubChem, Circular fingerprints) or shape/electrostatic similarity (ROCS, EON), could strengthen the prediction pipeline^37,38^. A consensus strategy combining these techniques may improve the robustness and predictive power.

From a structure-based perspective, this workflow does not incorporate docking due to the flexibility of the fascin binding site. However, for other targets or alternate conformations of fascin, it would be feasible to include structure-based methods, such as blind docking, standard docking, or molecular dynamics simulations with MM/PBSA scoring. These enhancements would allow for a more comprehensive evaluation of ligand-target interactions, thereby improving the predictive accuracy and translational relevance of the workflow.

In summary, we developed an automated pipeline for the discovery of novel anticancer compounds using a known molecule as a reference. This tool enabled the identification of two novel compounds with improved bioactivity compared to monastrol. Beyond fascin, this approach holds promise for broader applications in anticancer drug discovery and may serve as a foundational methodology for exploring inhibitors of other cancer-relevant targets.

## Conclusion

By implementing an efficient Ligand-Based Virtual Screening workflow, this study led to the discovery of two novel anti-fascin agents from the extensive combinatorial library of Enamine. Unlike previous studies that focused on smaller libraries of FDA-approved compounds, our approach explored a broader chemical space. By using the pharmacophore features of monastrol, the model identified two new fascin inhibitors with unique chemical properties and potential freedom from intellectual property constraints. Furthermore, we provided molecular insights into the structural interactions of these compounds with fascin, indicating that this protocol could be applied to discovering additional anti-fascin agents. Finally, we described a novel pipeline that integrates *in silico* and experimental validations to identify promising anticancer compounds. This approach could be extended to other cancer targets in the future, and may contribute to drug discovery efforts against different classes of therapeutic targets.

## Supporting information

In vitro validation of binding to fascin1 by DSF and fascin bundling assays

## Data availability

No datasets were generated or analysed during the current study.

## Abbreviations

BD: Blind Docking
CRC: Colorectal cancer
EGFR: Epidermal growth factor receptor
GLUT1: Glucose Transporter 1
GUI: Graphical user interface
HTS: High-throughput screening
MD: Molecular Dynamics
SAC: Serrated colorectal adenocarcinoma
Tm: Melting Temperature

## Acknowledgments

This study was supported by a grant from the Scientific Foundation of the Spanish Association against Cancer (Predoctoral Fellowship). This research was funded by JUNTA ANDALUCIA GRANT, the Andalusian Regional Government, through Grant Proyectos de Excelencia (P18-RT-1193). Project support for this research was provided in part by the European Union Horizon 2020 Grant (REVERT project GA848098), Instituto de Salud Carlos III (PI23/00601), and FEDER/Junta de Andalucía-Consejería de Transformación Económica, Industria, Conocimiento y Universidades (PY20_00678 and B-BIO-18-UGR20). The authors acknowledge the computing resources and technical support provided by the Plataforma Andaluza de Bioinformática at the University of Málaga. Powered@NLHPC research was partially supported by the super-computing infrastructure of NLHPC (ECM-02). The author(s) gratefully acknowledge the computational resources provided by the Barcelona Supercomputing Center at MareNostrum5 under project BCV-2025-1-0004. This work was supported by Ministerio de Ciencia e Innovation through the grant Ramón and Cajal program (Grant RYC2022-037702-I, funded by MCIU/AEI/10.13039/501100011033 and by the ESF+)

## Declaration of generative AI and AI-assisted technologies in the writing process

During the preparation of this work the author(s) used ChatGPT and Paperpal: AI Academic Writing Tool in order to improve language and readability. After using this tool/service, the author(s) reviewed and edited the content as needed and take(s) full responsibility for the content of the publication.

## Author contribution

**Alejandro Rodríguez-Martínez**: Conceptualization, Methodology, Software, Validation, Formal analysis, Investigation, Data Curation, Writing - Original Draft, Writing - Review & Editing, Visualization. **Lucía Giraldo-Ruiz**: Validation, Investigation, Writing - Original Draft, Writing - Review & Editing, Visualization. **Maria C. Ramos**: Investigation, Resources, Writing - Original Draft, Writing - Review & Editing, Supervision, Project administration, Funding acquisition. **Irene Luque**: Methodology, Formal analysis, Investigation, Resources, Writing - Original Draft, Writing - Review & Editing, Supervision, Project administration, Funding acquisition. **Diogo Ribeiro**: Validation, Investigation, Writing - Original Draft, Writing - Review & Editing, Visualization. **Fátima Postigo-Corrales**: Validation, Investigation, Writing - Original Draft, Writing - Review & Editing, Visualization. **Begoña Alburquerque-González**: Investigation, Writing - Original Draft, Writing - Review & Editing. **Silvia Montoro-García**: Methodology, Investigation, Resources, Writing - Original Draft, Writing - Review & Editing. **Ana Belén Arroyo-Rodríguez**: Validation, Investigation, Writing - Original Draft, Writing - Review & Editing. **Pablo Conesa-Zamora**: Methodology, Formal analysis, Investigation, Resources, Writing - Original Draft, Writing - Review & Editing, Supervision, Project administration, Funding acquisition. **Ana María Hurtado**: Validation, Investigation, Writing - Original Draft, Writing - Review & Editing. **Ginés Luengo-Gil**: Methodology, Formal analysis, Investigation, Resources, Writing - Original Draft, Writing - Review & Editing, Visualization, Supervision, Project administration, Funding acquisition. **Horacio Pérez-Sánchez**: Conceptualization, Software, Formal analysis, Investigation, Resources, Writing - Original Draft, Writing - Review & Editing, Supervision, Project administration, Funding acquisition.

