## Supplementary material for "Characterizing Compounds Targeting Colorectal Cancer Derived From Monastrol Using High-Through Screening of an Extensive Combinatorial Library": In vitro validation of binding to fascin1 by DSF and fascin bundling assays

[1] Structural Bioinformatics and High Performance Computing Research Group (BIO-HPC), HiTech Innovation Hub, UCAM Universidad Católica de Murcia, Murcia 30107, Spain; Health Sciences PhD Program, Universidad Católica deMurcia UCAM, Murcia 30107, Spain

[2] Structural Bioinformatics and High Performance Computing Research Group (BIO-HPC), UCAM Universidad Católica de Murcia (UCAM), 30107 Murcia, Spain

[3] Departamento de Química Física, Instituto de Biotecnología y Unidad de Excelencia de Química aplicada a Biomedicina y Medioambiente Universidad de Granada. Campus Fuentenueva s/n 18071 Granada, Spain

[4] Genetics, Molecular Pathology and Rare Diseases Research Group, Universidad Católica de Murcia (UCAM); Guadalupe, 30107, Spain

[5] Molecular Pathology and Pharmacogenetics Research Group, Instituto Murciano de Investigación Biosanitaria (IMIB), Hospital General Universitario Santa Lucía; Cartagena, 30202, Spain

[6] Health Sciences PhD Program, Universidad Católica de Murcia UCAM, Campus de los Jerónimos nº135, Guadalupe, Murcia, 30107, Spain

### *In vitro validation of binding to fascin1 by DSF and fascin bundling assays*

The effect of Z118298144 on fascin bundling activity was evaluated using both the DSF technique and the fascin bundling imaging-based assay. In the high-content imaging assay, compound Z118298144 induces only a partial or moderate reduction in the formation of F-actin bundles, suggesting a less potent inhibitory activity compared to stronger fascin inhibitors. Nevertheless, the compound produced a clear thermal shift in the Differential Scanning Fluorimetry (DSF) assay, supporting a significant structural interaction with fascin (Figure S1). This is consistent with the moderate activity observed in our HTS assays, where the presence of Z118298144 resulted in less structured F-actin assemblies. Overall, Z118298144 appears to engage structurally with fascin (as shown by DSF), but with a more moderate functional impact in cellular contexts.

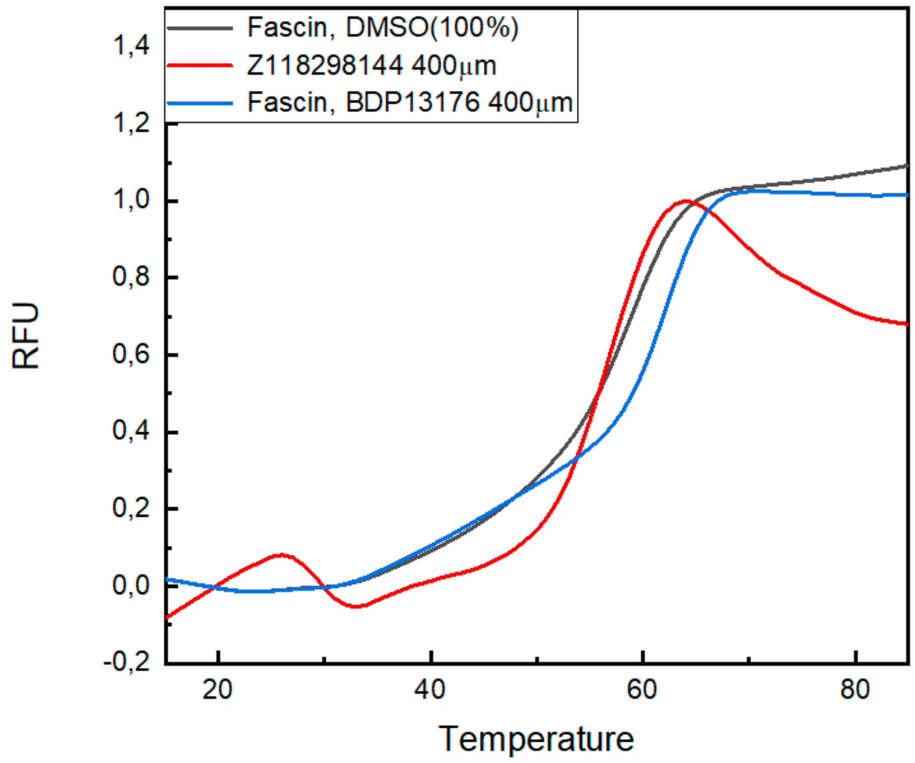


**Figure S1.** Differential Scanning Fluorimetry (DSF) analysis of Fascin stability in HEPES-NaCl buffer under three experimental conditions. The black trace represents the control condition (1 µM Fascin with 100% DMSO). The blue trace corresponds to Fascin (1 µM) in the presence of compound BDP13176 at 400 µM, while the red trace shows the effect of compound Z118298144 at the same concentration. The addition of Z118298144 results in a moderated destabilization of Fascin, as evidenced by the absence of a canonical thermal unfolding curve and alteration in the fluorescence signal. All experiments were conducted in 20 mM HEPES, 100 mM NaCl, pH 7.35.

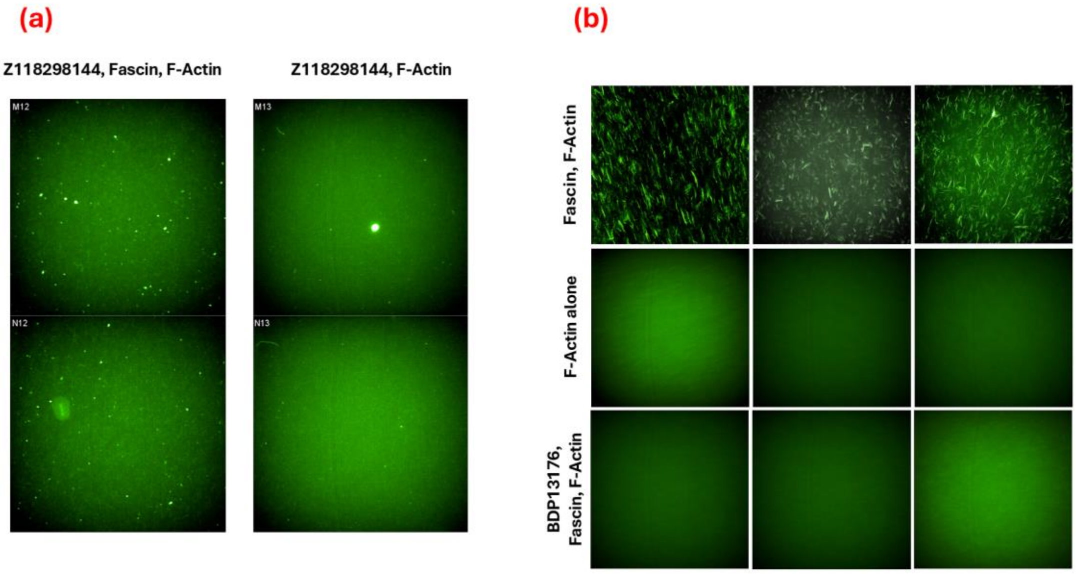


**Figure S2.** Representative fluorescence images from a high-content imaging assay evaluating Fascin-dependent F-actin bundling. The condition with actin and Z118298144 (0.5 µM actin + 400 µM compound) shows an absence of F-actin bundles, indicating that the compound does not directly interfere with actin polymerization and suggesting specificity for Fascin inhibition. Control assays include a negative control (0.5 µM Fascin + 0.5 µM actin + 4% DMSO), where robust F-actin bundles are visible, confirming Fascin's bundling activity; an actin-only condition (0.5 µM), which shows no fiber formation, indicating that actin does not self-assemble under these conditions; and a positive control (1 µM Fascin + 0.5 µM actin + 400 µM BDP13176), in which F-actin bundling is completely disrupted due to effective Fascin inhibition by BDP13176.

On the other hand, compound Z17544625 did not yield positive results in these types of assays. Nevertheless, it was also evaluated in subsequent cell-based experiments, where it demonstrated activity in terms of cancer cell viability and migration.
